# Structures of topoisomerase V in complex with DNA reveal unusual DNA binding mode and novel relaxation mechanism

**DOI:** 10.1101/2021.08.20.457074

**Authors:** Amy Osterman, Alfonso Mondragon

## Abstract

Topoisomerase V is a unique topoisomerase that combines DNA repair and topoisomerase activities. The enzyme has an unusual arrangement, with a small topoisomerase domain followed by 12 tandem (HhH)_2_ domains, which include three AP lyase repair domains. The unusual architecture of this enzyme bears no resemblance to any other known topoisomerase. Here we present structures of topoisomerase V in complex with DNA. The structures show that the (HhH)_2_ domains wrap around the DNA and in this manner appear to act as a processivity factor. There is a conformational change in the protein to expose the topoisomerase active site. The DNA bends sharply to enter the active site, which melts the DNA and probably facilitates relaxation. The structures show a DNA binding mode not observed before and provide information on the way this unusual topoisomerase relaxes DNA.

## INTRODUCTION

The topological state of DNA in cells is regulated by the action of DNA topoisomerases (Bush, Evans-Roberts, & Maxwell, 2015; Corbett & Berger, 2004; Pommier, Sun, Huang, & Nitiss, 2016; J. C. Wang, 2002). Different cellular process, such as transcription and recombination, can alter the topology of DNA and the action of topoisomerases helps maintain the correct topological state. In order to alter the topology of DNA, topoisomerases transiently break either one or two strand of DNA in order to allow movement of strands before the breaks are resealed. In this manner, topoisomerases can relax and supercoil DNA, catenate/decatenate, and knot/unknot DNA molecules, and in some cases RNA as well (Ahmad et al., 2016; H. Wang, Di Gate, & Seeman, 1996). Due to their involvement in crucial cellular processes, topoisomerases are the target of important chemotherapeutic agents (Pommier, Leo, Zhang, & Marchand, 2010).

Topoisomerases are found in all three domains of life, but the diversity in structure and sequence suggest that the major subtypes evolved independently (Forterre, Gribaldo, Gadelle, & Serre, 2007). All topoisomerases have in common the use of phosphotyrosine intermediates for transient cleavage through a transesterification mechanism (Corbett & Berger, 2004). Topoisomerases are classified into two types (I and II) based on whether they cleave one or two strands of DNA in a concerted manner. Within each type, topoisomerases are subclassified based on similarities at the sequence and structural levels. Type I enzymes, which cleave one DNA strand, are subclassified into three subtypes, IA, IB, and IC. Type IA enzymes are found in bacteria, archaea, and eukarya and employ an enzyme-bridged strand passage mechanism and share a strand passage structural domain with a very characteristic toroidal shape (Lima, Wang, & Mondragon, 1994). Type IB enzymes are also found in all domains of life, employ a swiveling or controlled rotation mechanism, and show clear structural and sequence similarities, despite some of them being much larger than other members of the same subtype (Corbett & Berger, 2004). The structure and mechanism of type IA and IB topoisomerases and their cellular role is well studied, and in both subtypes the general steps of the DNA binding and cleavage/religation mechanism are well understood.

The third type I topoisomerase subtype, IC, has only been found in the archaeal *Methanopyrus* genus (Forterre, 2006) and relaxes DNA using a controlled rotation/swiveling mechanism (Taneja, Schnurr, Slesarev, Marko, & Mondragon, 2007). Its only member is topoisomerase V, which was originally described as a type IB enzyme (Slesarev et al., 1993), but later classified into its own subtype due to the lack of sequence or structural similarities with other topoisomerases (Forterre, 2006; Taneja, Patel, Slesarev, & Mondragon, 2006). The exact cellular role of topoisomerase V in the hyperthermophile *Methanopyrus kandleri* is not known. Topoisomerase V works not only at extremely high temperatures (65°C to 122°C), but also in high salt concentrations (Slesarev, Lake, Stetter, Gellert, & Kozyavkin, 1994). It is also an unusual protein as it combines DNA repair and topoisomerization activities in the same polypeptide (Belova et al., 2001; Rajan, Osterman, & Mondragon, 2016; Rajan, Prasad, Taneja, Wilson, & Mondragon, 2013), a unique feature of this enzyme. Topoisomerase V is formed by a small, ∼30 kDa topoisomerase domain followed by twenty-four tandem Helix-hairpin-helix (HhH) repeats (Doherty, Serpell, & Ponting, 1996) arranged as twelve (HhH)_2_ domains (Belova, Prasad, Nazimov, Wilson, & Slesarev, 2002; Shao & Grishin, 2000). HhH repeats are associated with DNA lyase repair activity (Doherty et al., 1996; Shao & Grishin, 2000) and three of the twelve (HhH)_2_ domains have apurinic/apyrimidinic (AP) lyase and deoxyribose-5-phosphate (dRP) lyase activities (Rajan et al., 2016; Rajan et al., 2013). These twelve (HhH)_2_ domains also increase the topoisomerase activity as mutant topoisomerase V proteins with fewer repeats show decreasing relaxation activity (Belova et al., 2002; Rajan, Taneja, & Mondragon, 2010). (HhH)_2_ domains are usually found as isolated domains, not in tandem arrangements (Shao & Grishin, 2000), which is another unusual characteristic of topoisomerase V. The topoisomerase domain has no sequence or structural similarities to other topoisomerases or any other proteins, making it a unique fold (Forterre, 2006; Taneja et al., 2006). The active site residues have been identified (Rajan, Osterman, Gast, & Mondragon, 2014), but their three dimensional arrangement suggest a different catalytic mechanism from other topoisomerases (Rajan et al., 2014). In the structures of the free protein, the topoisomerase domain active site is inaccessible as it is covered by the (HhH)_2_ domains (Rajan et al., 2016; Rajan et al., 2013; Rajan et al., 2010; Taneja et al., 2006).

The way that topoisomerase V interacts with DNA, relaxes it, and recognizes lesions is not known. This stands in contrast to all other topoisomerase types, either I or II, where structures of complexes of the proteins with DNA are known (Bush et al., 2015; Corbett & Berger, 2004). The absence of structural information on topoisomerase V in complex with DNA has hampered efforts to understand fully not only its mechanism of DNA relaxation, but also the way AP/dRP lyase and topoisomerase activities are coupled and the conformational changes needed to expose the active site and bind DNA. Here, we present structures of topoisomerase V in complex with DNA containing an abasic site. The structures show the way the (HhH)_2_ domains embrace DNA and change conformation to expose the topoisomerase active site and accept DNA into it. Despite type IB and IC enzymes both using a controlled rotation DNA relaxation mechanism (Koster, Croquette, Dekker, Shuman, & Dekker, 2005; Taneja et al., 2007), the structures show that there are significant and substantial differences. The structures also suggest that the (HhH)_2_ domains are not only involved in DNA repair but may also serve as a processivity factor by encircling the DNA. Overall, the structures demonstrate a highly unusual DNA/protein complex that helps explain many of the features that make topoisomerase V a unique protein. The structures bring our understanding of type IC enzymes to a similar level as for all other topoisomerases.

## RESULTS

### Crystallization of complexes of topoisomerase V with DNA

Topoisomerase V is a non-sequence specific DNA binding protein that has AP lyase activity and interacts with DNA abasic sites (Rajan et al., 2016; Rajan et al., 2013). For this reason, co-crystallization trials were done using DNA fragments with abasic sites with the goal to target topoisomerase V to this position and create a more homogeneous complex. In addition, a 97 kDa fragment of Topoisomerase V mutated to have only a single AP lyase site (hereafter Topo-97(ΔRS2), see **Materials and Methods**) (Rajan et al., 2016) was used in all crystallization trials. After trying several DNA sequences and lengths, a 38 base pair (bp) DNA oligonucleotide with a single abasic site crystallized (**Materials and Methods**); the crystals were optimized by adding two complementary overhanging base pairs at the ends of the DNA. Once the structure of this complex was solved, it was evident that two protein monomers were bound to each DNA fragment, that the complementary overhangs were not interacting with each other, and that each protein monomer interacts with two different DNA molecules. Furthermore, the complexes were in the closed conformation of the protein, where the topoisomerase active site is buried.

The 38 bp DNA oligonucleotide was re-designed by making it symmetric from the center, with two abasic sites, and still containing overhangs (total length per strand 40 nucleotides) (**Materials and Methods**). The symmetric oligonucleotide led to a different crystal form. The structure of the symmetric oligonucleotide complex shows that each DNA fragments binds two protein molecules, but in the open conformation and with the topoisomerase active site accessible. The two proteins are related by a two-fold axis, but this axis does not coincide with the DNA sequence two-fold axis, which lies between two base pairs. Instead, the two-fold axis passes through a base pair thus creating an asymmetry in the DNA structure. A second symmetric DNA molecule (39 bp long) was designed with the two-fold axis passing through a central base pair. This last construct produced a structure with a symmetric DNA molecule and with the protein and DNA two-fold axes coinciding. In addition, a third and slightly longer DNA molecule (40 bp) based on the 38 bp symmetric oligonucleotide was tried, it produced slightly better diffraction. Overall, four complexes were crystallized and solved (**Materials and Methods**). Three of them are closely related and are formed by symmetric DNA with two abasic sites each; the fourth one is asymmetric and contains a single abasic site.

### Overall structure of topoisomerase V with asymmetric DNA

Topo-97(ΔRS2) consists of the topoisomerase domain at the N-terminal end and 10 (HhH)_2_ domains arranged in tandem (**Figure 1A**). The crystals of the complex formed by Topo-97(ΔRS2) with a 38 bp asymmetric DNA oligonucleotide containing one abasic site diffract to a 3.24 Å in the best direction but only to ∼3.9 Å in the other directions (**Table II**). This anisotropy resulted in an electron density map of uneven quality and although diffraction extends to higher resolution, the map mostly corresponds to a medium resolution structure. Nevertheless, the placement of the protein molecules was unambiguous and a previously unseen region, repeat 7 and Linker helix II, linking repeats 7 and 8, could be seen clearly (**Figure 1**). The region was built with the aid of the structure from the symmetric complexes, which diffract to higher resolution. The DNA molecules were easily recognized and built. The asymmetric unit of the crystal contains two complexes, each formed by a Topo-97(ΔRS2) protein molecule and two different DNA molecules. This arrangement is unusual, with each protein monomer bound to only half of each DNA molecule. The center of two of the half-bound DNA molecules sits on crystallographic two fold axes, recreating the full length DNA (**Figure 1 – Figure Supplement 1**). The other DNA molecule is bound by the two protein monomers. The two complexes in the asymmetric unit are very similar, but not identical (rmsd between Cα atoms ∼ 1.0 Å).

**Figure 1.**
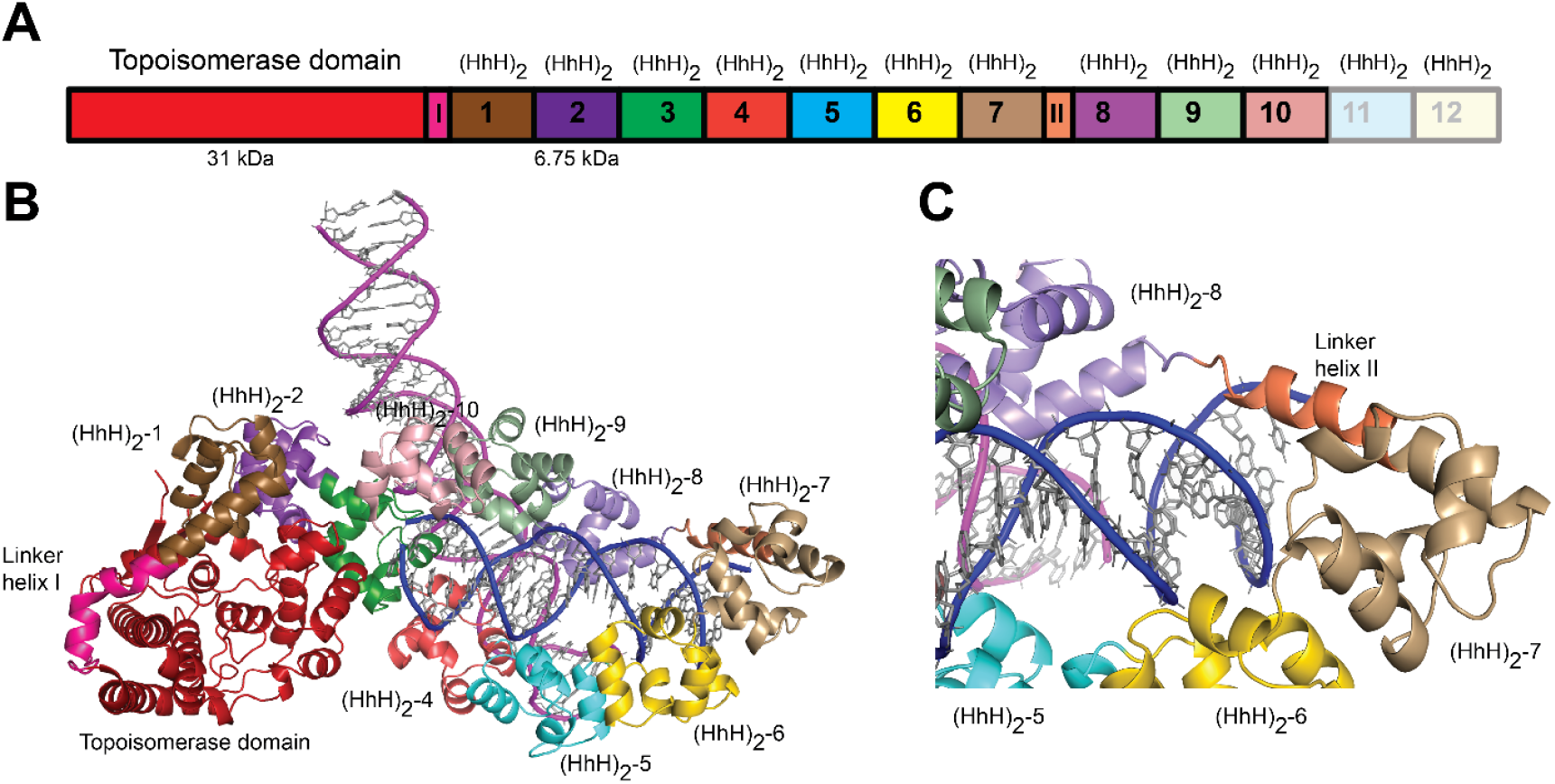
Structure of topoisomerase V in complex with asymmetric DNA. **A)** Schematic diagram of the domain organization of topoisomerase V. The protein contains a small, 31 kDa topoisomerase domain followed by twelve (HhH)_2_ domains each formed by two (HhH) repeats. There are linker helices between the topoisomerase and the first (HhH)_2_ domain (LI) as well as between repeats 7 and 8 (LII). Repeats 11 and 12 are not part of the structure. **B)** Cartoon showing the structure of topoisomerase V in complex with asymmetric DNA. Each protein monomer binds two half DNA molecules. The topoisomerase domain remains blocked by the (HhH)_2_ domains preventing access to the active site. **C)** Close up view of the region around repeats 7 and 8, which are connected by Linker helix II. The linker helix serves to connect two sets of domains and sits above the major groove of the DNA. In this structure, one DNA is surrounded by the (HhH)_2_ domains while the other sits between (HhH)_2_ domains. The domains are colored using the same scheme as in the **A** diagram.

Each complex shows the protein in the closed conformation with the (HhH)_2_ domains surrounding one DNA molecule. The second DNA molecule interacts with a subset of (HhH)_2_ domains in an unanticipated manner (**Figure 1B**). The two DNA molecules bind the protein almost perpendicularly, at a ∼120º angle (**Figure 1B**). In previous structures of fragments of topoisomerase V (Rajan et al., 2016; Rajan et al., 2013), the sixth and seventh (HhH)_2_ domains and the linker joining it to the eighth one were partially or fully disordered. In the complex structure this region is ordered in one of the monomers and shows that the seventh repeat forms an (HhH)_2_ domain followed by a long helix (Linker helix II) that connects it to the eighth (HhH)_2_ domain. This long helix sits in the major groove of one DNA molecule, making the (HhH)_2_ domains wrap around the DNA (**Figure 1C**). Aside from the ordering of this region and the swiveling of the last three (HhH)_2_ domains, no other major changes are seen in the protein with respect to the structure of the same fragment in the absence of DNA (Rajan et al., 2016). Thus, in the closed conformation the main change in the protein structure is the movement of (HhH)_2_ repeats 8-10 to enclose the DNA and the ordering of the seventh domain and the linker region.

The protein binds the two DNA molecules through the (HhH)_2_ domains. One of the DNA molecules is almost surrounded by the (HhH)_2_ domains with one end abutted against the topoisomerase domain (**Figure 1B** and **Figure 1 – Figure Supplement 1**). As the enzyme is in the closed conformation, the DNA cannot interact with the topoisomerase domain or enter the active site. The second DNA molecule sits in a positively charged groove formed by the (HhH)_2_ domains. The two DNA molecules come close together at this point but do not interact directly. The DNA molecules are in the B-conformation with one of them bent, but mostly having canonical DNA parameters. Interestingly, the same DNA sequence without the abasic site also crystallized under the same conditions, suggesting that the presence of the DNA abasic site had no effect on the binding of the protein or the conformation of the DNA. Not surprisingly, the DNA abasic site was not apparent in the structure and does not appear to cause any deviations from canonical B-DNA.

### Overall structure of topoisomerase V with symmetric DNA

The use of a symmetric DNA with two abasic sites resulted in a new crystal form with one DNA molecule bound by two Topo-97(ΔRS2) monomers (**Figure 2 – Figure Supplement 1)**. Different lengths of DNA were tried to improve the crystal quality, but all of them suffered from anisotropic diffraction. The best diffraction was from crystals with a 40 bp DNA oligonucleotide with 2 base overhangs at each end (**Materials and Methods**). These crystals diffract to 2.92 Å in the best direction but only to ∼3.5 Å in the other directions (**Table III**). Crystals with 39 or 40 bp DNA and including 2 base overhangs were also anisotropic, but served to provide information on the path of the DNA.

In the structure, one DNA molecule is surrounded by two protein monomers through the (HhH)_2_ domains (**Figure 2A, 2C**). The two proteins in the asymmetric unit are very similar in conformation and are related by an almost perfect non-crystallographic two-fold axis (rmsd between Cα ∼ 1.0 Å). For this reason, only one monomer is described hereafter. The (HhH)_2_ domains can be divided into two subsets, (HhH)_2_ domains 1 to 7 and 8 – 10. Each (HhH)_2_ domains subset has the same conformation as in the DNA-free protein, showing that they move as rigid groups. Similarly to the asymmetric DNA complex, the (HhH)_2_ domains change direction after the seventh domain and the helix linking domains seventh and eighth is well-ordered. The two subsets form a loop-like structure where the turn is formed by the helix connecting domains 7 and 8. The topoisomerase domain is shifted with respect to the (HhH)_2_ domains and is accessible to the DNA. The topoisomerase domain moves as a rigid body with the only changes observed confined to the helix linking the topoisomerase domain and the first (HhH)_2_ domain(Linker helix I) (**Figure 2B**). This helix, which in the DNA-free structure is an extension of the first helix of the (HhH)_2_ domain, is broken and in this way separates the topoisomerase domain from the (HhH)_2_ domains (**Figure 2 – Figure Supplement 2**). The changes in this helix are the only major changes observed when the DNA-free and DNA-bound structures are compared. The movement of the topoisomerase domain exposes the active site (**Figure 2**), which in all other structures was inaccessible, and allows the end of the DNA molecule to enter the topoisomerase domain and interacts with residues near the active site region (**Figure 2D**).

**Figure 2.**
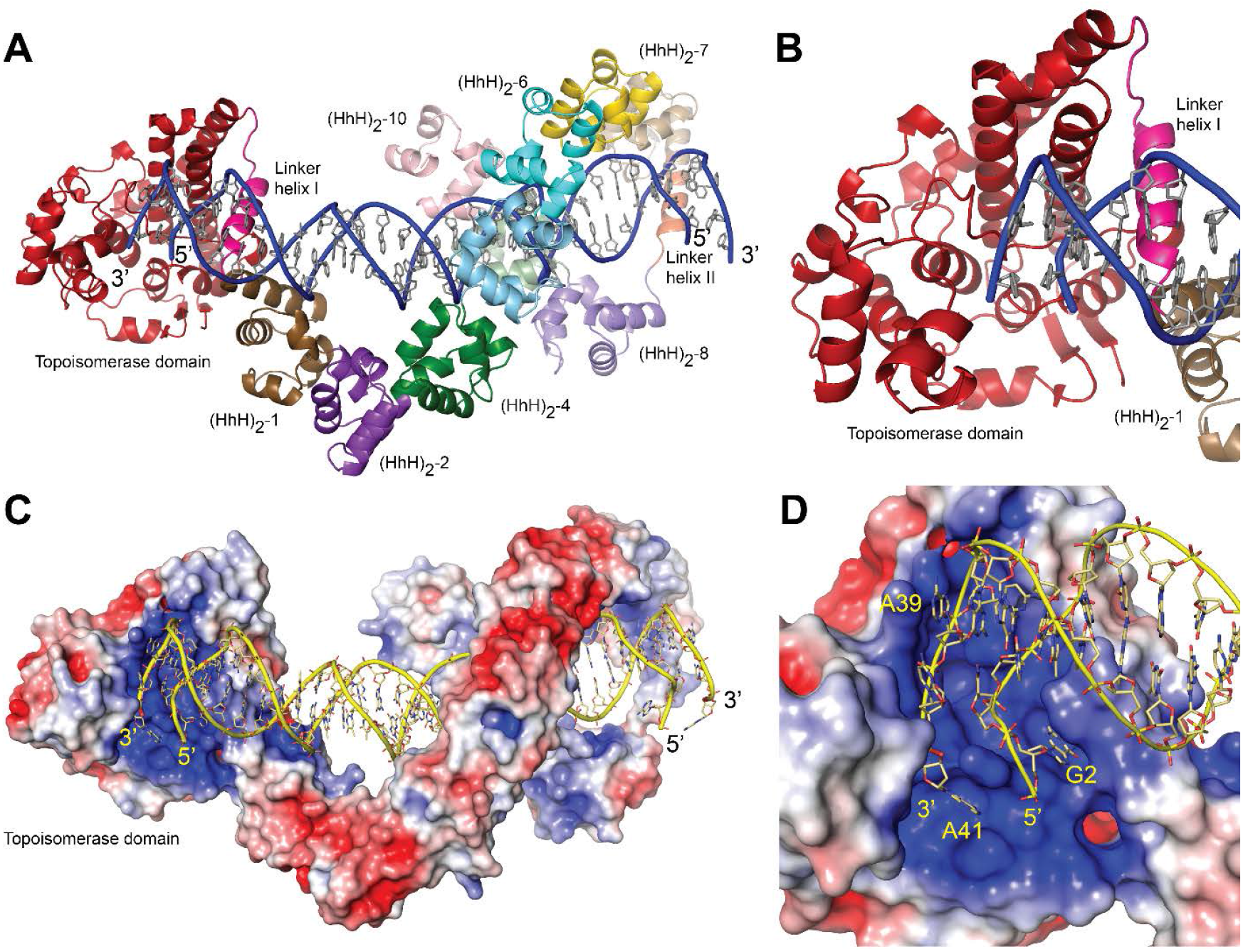
Structure of topoisomerase V in complex with symmetric DNA. **A)** Cartoon showing the structure of topoisomerase V in complex with symmetric 40 bp DNA. Each DNA molecule binds two protein molecules in a symmetric manner; only one protein molecule is shown. In the structure, the topoisomerase domain moves away from the (HhH)_2_ domains allowing access to the active site. The (HhH)_2_ domains wrap around the DNA. **B)** Close up view of the topoisomerase domain (red). Linker helix I (pink) changes conformation to allow movement of the domains and expose the active site. **C)** Electrostatic surface of topoisomerase V in the bound conformation. The interior of the cavity formed by the (HhH)_2_ domains is slightly positively charged, forming a region where the DNA can bind. The end of the DNA molecule enters the topoisomerase active site. **D)** Close up view of the topoisomerase domain active site. The DNA enters a highly positively charged region where the active site residues are located. To enter the active site, the DNA bends and the base pairing between bases is broken. The protein domains are colored as in Figure 1. The surface is colored with a blue to red gradient from +5 to −5 K_b_T/e_c_.

The structure with a 38 bp oligonucleotide revealed that the non-crystallographic two-fold axis passes through a base pair of the DNA molecule, making the 38 and 40 bp molecules sit asymmetrically in the complex; one half comprises 18 or 19 bp and the other half 19 or 20 bp for the 38 and 40 bp DNA molecules, respectively. This asymmetry translates in an asymmetry in the path of the DNA, even though the protein monomers are identical. To regularize the complex, a DNA molecule with an odd number of base pairs (39 bp) was used. In this case, the two-fold axis passes through the central base pair, the two halves of the DNA molecule are identical, and the path of the DNA is symmetric. The even more symmetric structure did not show any differences in the proteins.

### Structure of the DNA in the complexes

In the asymmetric complex each protein monomer interacts with two different DNA molecules. One of them is slightly bent and is shared by two proteins, while the second one is straighter. In either case, it was not possible to observe the abasic site. Unlike the asymmetric complex, the DNA in the symmetric complex shows clearly the presence of the abasic site. The structure shows that the abasic site is accommodated by unstacking the complementary base, which moves to lie in the major groove of the DNA (**Figure 3**). The sugar of the abasic site remains in the expected position. The result of the rearrangements is that the adjacent base to the abasic site moves to occupy the vacant space left by the abasic site. These movements creates a kink in the DNA leading to an opening of the major groove and a narrowing of the major groove just before the abasic site.

**Figure 3.**
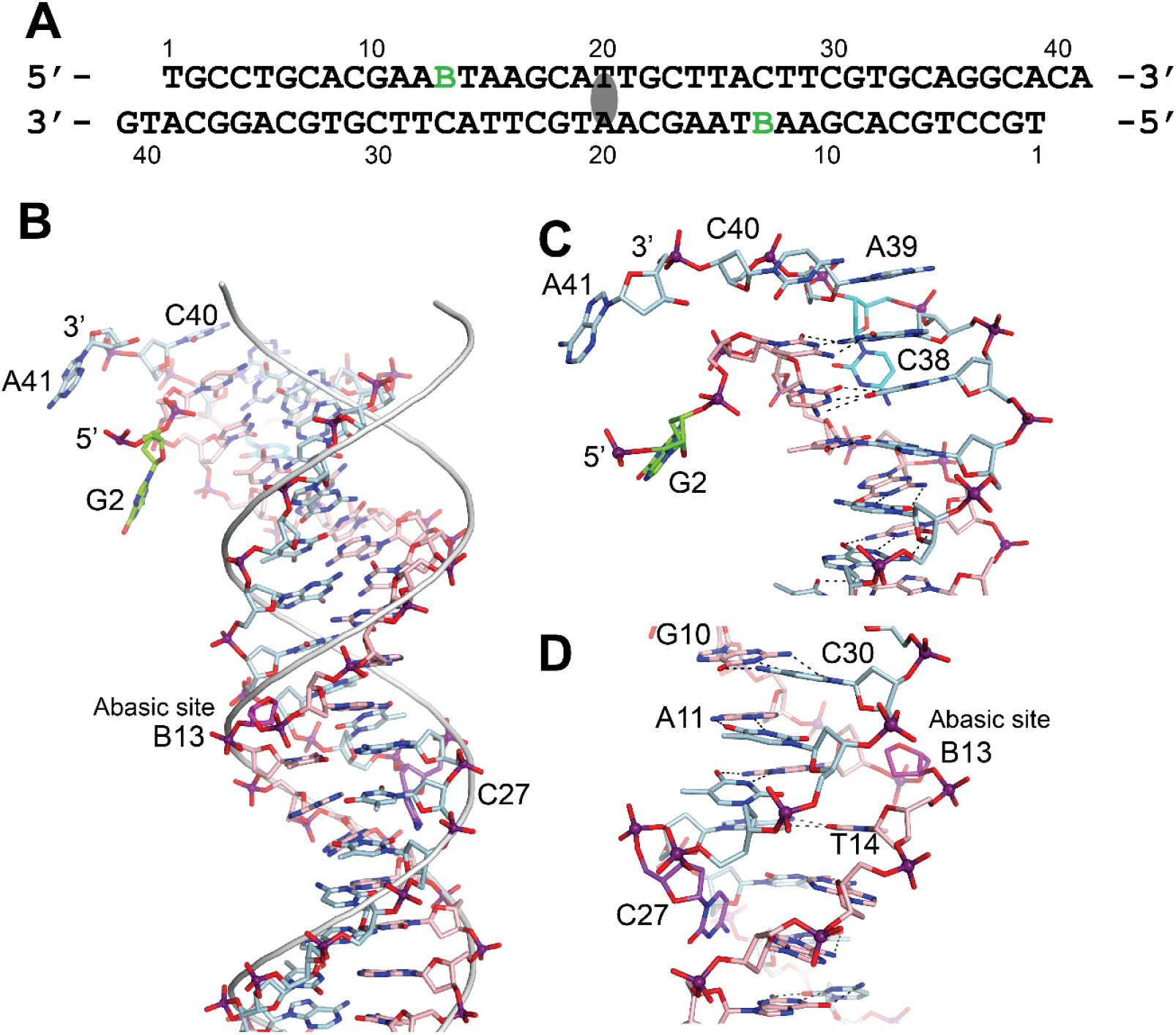
DNA in the structure of topoisomerase V with 39 bp DNA. **A)** Sequence of the 39 bp DNA in the structure. The DNA is symmetric, with the two-fold axis passing through the central base pair. The abasic sites are shown with a green B. **B)** Stick diagram of one half of the 39 bp DNA in the complex. The DNA molecule is bent due to the presence of the abasic sites. For comparison, the phosphate backbone path of a B DNA molecule is shown as a grey tube. Note that the bending occurs around position B13, the abasic site. For clarity, the carbon atoms in one strand are colored in light blue and on the complementary strand in pink. **C)** Close up view of the end of the molecule. The DNA is bent where it enters the active site and the bending causes the base paring to break. The second nucleotide, G2 (green), is not base paired to its corresponding base pair, C38 (cyan). The latter unstacks from the helix to allow the next nucleotide A39, to interact with the protein. The first nucleotide, T1, is disordered in the structure. **D)** Close up view of the region around the abasic site. The abasic site (B13, magenta), causes the DNA to bend. The corresponding base pair, C27 (purple), completely unstacks from the helix and the base enters the minor groove. The unstacking of C27 plus the movement of the sugar in the abasic site allows the stacking of bases to continue without any gaps, despite the presence of the abasic site.

The bending around the abasic site allows the DNA to approach the active site (**Figure 2**). In addition, in order to interact with the protein the DNA is very sharply bent where it enters the active site. This sharp bend makes the two DNA strands melt and the last two complementary nucleotides are not base paired (T1:A39,G2:C38) (**Figure 3**). Instead, C38 is unstacked and enters the minor groove, while A39 stacks on top of G37 (**Figure 3**). The melting allows the last four nucleotides on the 3’ end of the DNA to enter the protein and do not interact with the other DNA strand (**Figure 3**). The 5’ end of the DNA has two unpaired nucleotides, the first one mostly disordered as it has moved away from the protein whereas the second nucleotide, G2, enters a pocket in the protein, stacked against the side chains of Tyr289 and Arg109.

### DNA-protein interactions

The (HhH)_2_ domains surround most of the DNA coming in close contact with the phosphate backbone at many points. They form a loop with a positively charged interior that accommodated the DNA, but appears to have few close contacts (**Figure 2**). There are no contacts of the (HhH)_2_ domains with the bases, only with the backbone, which is not unusual for a non-sequence specific DNA binding protein. Repeats 3-6 and repeat 9 make contacts with the phosphate backbone while the rest of the repeats only surround it, but do not come close to it. Interestingly, even though many of the repeats do not contact the DNA directly, they have positively charged residues facing the DNA, creating an overall positively charged environment around the DNA (**Figure 2**).

The abasic sites are not in direct contact with the protein even though the abasic sites were introduced as a possible target for the single intact repair (HhH)_2_ domain. Repeat 6 contains an AP/dRP lyase active site that includes lysines 566, 570, and 571; mutations of any of these three residues is deleterious for activity (Rajan et al., 2013). In the complex structure, lysines 570 and 571 face the DNA phosphate backbone, but are not close enough to contact it. Lysine 566 faces away from the DNA and cannot contact the DNA. This suggest that either lysine 570 or 571 are the likely nucleophile, but it is not clear how they would recognize or cleave the abasic site. The abasic site is in close contact to repeat 9, but this repeat has not been implicated in AP/dRP lyase activity (Rajan et al., 2013). Thus, it is not clear from the structures how the protein recognizes the abasic site and cleaves it.

The active site is exposed in the structure and reveals a highly positively charge region where the DNA enters. As mentioned above, the DNA in this region melts due to a sharp bend. This groove is narrow but expands on the side that faces the solvent and where the DNA exits. Binding of DNA in this region is plastic. Oligonucleotides of different lengths can be accommodated by following a slightly different path before entering the active site (**Figure 3 – Figure Supplement 1**). The 5’ end of one strand always follows the same path whereas the 3’ end does not. The last nucleotide on the 3’ end always ends up near the active site, but the unpaired nucleotides before the last one can follow slightly different paths. The contacts between the protein and the DNA are mostly with the phosphate backbone, with the bases facing the solvent, although one nucleobase at the 5’ end is tightly wedged between two protein side chains. The wedging of this nucleobase may help keep the non-cleaved strand in position during swiveling.

### Active Site

The residues forming the active site were identified from the structure of a 61 kDa fragment (Taneja et al., 2006) and later confirmed by site directed mutagenesis (Rajan et al., 2014). Aside from Tyr226, the active site tyrosine, five residues were identified as playing a role in the cleavage/religation reaction: Arg131, Arg144, His200, Glu215, and Lys218. The structure of the complex shows that aside from Lys218 all these residues are in the vicinity of the DNA (**Figure 4**). Based on the structure, the putative scissile phosphate, termed here P0, would correspond to the last phosphate in the oligonucleotide but is not present in the structures, as the 3’ end of the oligonucleotides is dephosphorylated. In all oligonucleotides studied, the 3’ end of the last nucleotide is in the same general location, near the tyrosine, but pointing away from it. A simple rotation around one of the bonds would bring it near the active site tyrosine. As the protein opens to the solvent in this region, there are few obstacles to hinder the rotation and allow the phosphate backbone to enter completely into the active site; the nucleobase could easily move into the solvent region. Arg131 and Arg144 both contact the P-1 phosphate, the phosphate group immediately 5’ of the scissile one (**Figure 4**). Their orientation is such that they could also contact the P0 phosphate if it was present as they are in the region between the two phosphates. His200 and Glu215 are hydrogen bonded to each other. In the complex structure, they are too far to contact the DNA directly, but Glu215 could contact the P0 phosphate with minimal side chain rearrangements. Finally, Lys218 is not making contacts with the DNA but is close enough to Tyr226 that it could contact the P0 phosphate. Thus, all residues that have been involved in cleavage and religation are positioned in a manner consistent with their previously assigned roles (Rajan et al., 2014).

**Figure 4.**
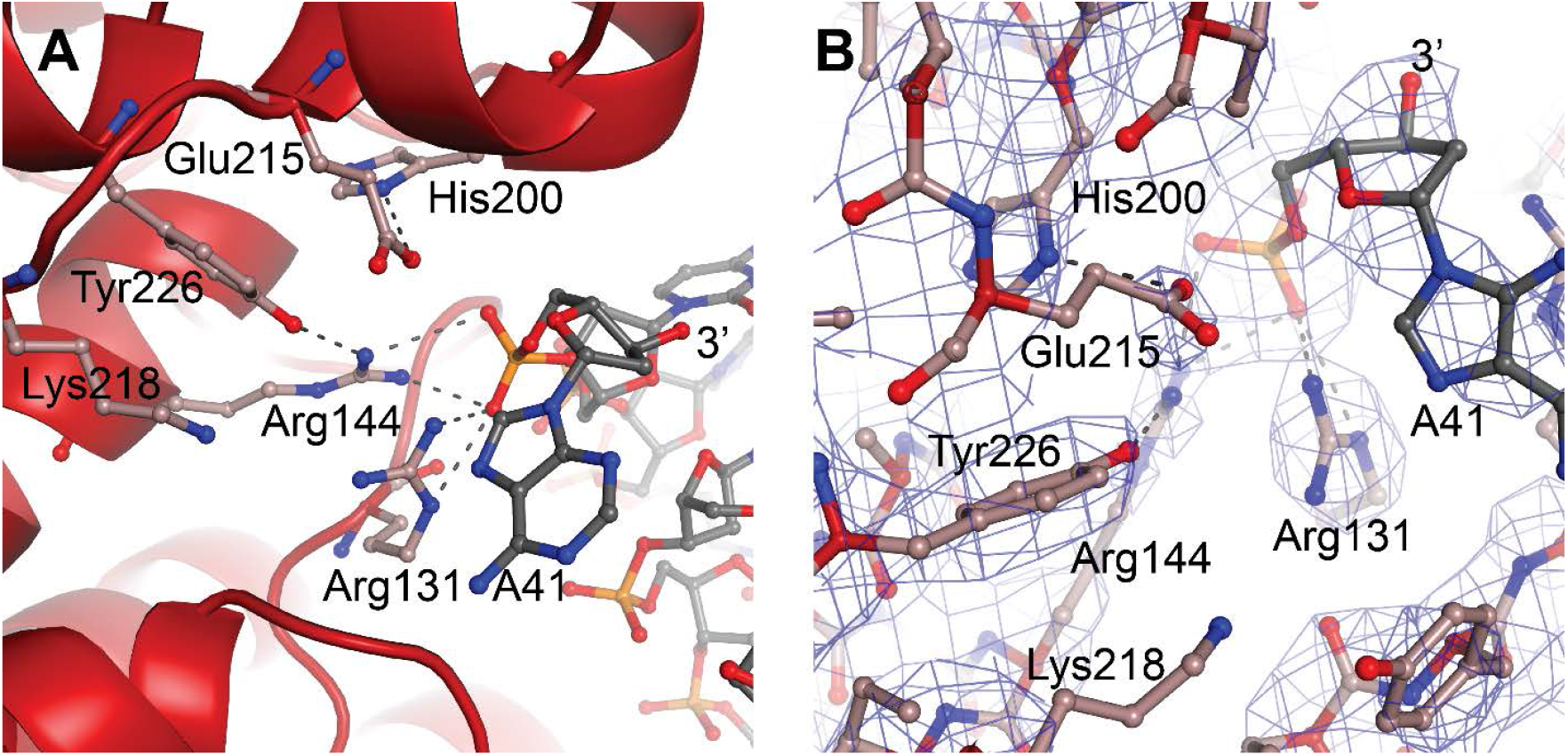
Topoisomerase V active site. **A)** Diagram of the active site in the topoisomerase V with 40 bp symmetric DNA complex. The diagram shows the side chains that have been implicated in the cleavage/religation reaction. The DNA approaches the active site and comes close to the active site tyrosine (Tyr226). The preceding phosphate (P1) is contacted by two arginines, Arg131, and Arg144. His200 is too far away from the phosphate backbone, but is hydrogen bonded to Glu215. The latter is in a position suitable to contact the DNA phosphate backbone Finally, Lys218 is in a position where it could contact the phosphate backbone during cleavage/religation. **B)** Simulated annealing omit map of the topoisomerase V with 40 bp symmetric DNA complex structure. The diagram presents another view of the active site as well as the omit electron density around that region at the 1 σ level.

It is not surprising that the only major change around the active site between the free protein and the protein/DNA complex is the movement of the HhH_2_ domains. At this resolution, it is not possible to discern large changes in conformation in any of the side chains around the active site tyrosine or in the nearby regions contacting the DNA. It appears that the active site is largely preassembled, even though in the closed conformation it is buried. It is likely that some changes may occur during cleavage and religation, but these changes do not have to be major.

## DISCUSSION

The general outlines of the DNA binding mode and relaxation mechanism is understood for many topoisomerases (Bush et al., 2015; Corbett & Berger, 2004), but this is not the case for type IC enzymes. In the latter case, the absence of information on the way they recognize and bind DNA has limited our understanding of the mechanism of this topoisomerase subtype. In addition, the lack of sequence or structural similarities between topoisomerase V and other topoisomerases meant that it was not possible to deduce the DNA binding mechanism based only on similarities. The presence of multiple (HhH)_2_ domains as well as biochemical information suggested that these domains were involved in DNA binding (Belova et al., 2002; Rajan et al., 2010), but it was not clear how tandem domains recognize and bind DNA. The structures presented here show the way topoisomerase V interacts with DNA, both through the (HhH)_2_ domains and the topoisomerase domain. DNA binding by the (HhH)_2_ domains is unusual and in a way not previously observed in other proteins. The tandem (HhH)_2_ domains wrap around the DNA forming a loop-like structure that encircles the DNA. Typically, HhH repeats are found as single repeats or forming (HhH)_2_ domains (Shao & Grishin, 2000), not as tandem arrangements. Some proteins that include (HhH)_2_ domains dimerize to have two (HhH)_2_ domains next to each other (for example (Nowotny & Gaur, 2016)), but the (HhH)_2_ domains are not arranged in a tandem configuration in the protein. For this reason, even though HhH repeats and (HhH)_2_ domains are well studied, it was not possible to model DNA binding by topoisomerase V based on known structures. Unlike other (HhH)_2_ domain-containing proteins, where the domains are involved in DNA binding and repair, in topoisomerase V some (HhH)_2_ domains surround the DNA but do not contact it, others contact it, and only three out of twelve are directly involved in repair. When comparing individual (HhH)_2_ domains with repair enzymes that interact with DNA it is clear that the interaction observed in topoisomerase V is different from the one observed in other (HhH)_2_ domain complexes with DNA. It is not possible to establish whether the topoisomerase V repeats directly involved in DNA repair will bind DNA similarly to the way other DNA repair enzymes with (HhH)_2_ do. The (HhH)_2_ domains that contain the AP/dRP lyase activity do not engage directly with the DNA abasic sites in the structure. For this reason, the question on how topoisomerase V recognizes DNA lesions is not completely answered by the current structure. To understand the way the enzyme recognizes and cleaves abasic sites additional structural information on the interactions of the repair domains and an abasic site is needed.

The overall DNA binding mode by topoisomerase V is unusual. The (HhH)_2_ domains surround the DNA loosely covering almost four helical turns. Topoisomerase V clamps around DNA by having two sets of tandem repeats of (HhH)_2_ domains that follow the path of DNA in opposite directions. The region between repeats 7 and 8 serves as the turning point to permit the repeats to change direction and encircle the DNA. The Topo-97(ΔRS2) fragment is missing the last two (HhH)_2_ domains plus a few amino acids of unknown structure at the C-terminus. Based on the structures described here, it is likely that the last two (HhH)_2_ domains continue the same path as the previous three and interact with the topoisomerase domain and completely enclose the DNA forming two full loops. Given the length of DNA covered, there are few direct interactions with the phosphate backbone and very few with the nucleobases. Instead, it appears that the (HhH)_2_ domains create an enclosed positively charged track or groove where the DNA can travel. Other proteins that use tandem repeats of DNA binding domains do not make a turn to change direction while binding DNA. Instead, they wrap around the DNA in a single direction. For example, zinc finger containing proteins (for instance Zif268 or Gli (Pavletich & Pabo, 1991, 1993)) and TAL effector proteins binding domains (Mak, Bradley, Cernadas, Bogdanove, & Stoddard, 2012) interact with the DNA by having tandem repeats that wrap around the DNA, but they extend along the DNA, not turning to form a loop-like structure that surrounds it.

DNA binding by topoisomerase V does not require all (HhH)_2_ domains, but its activity is enhanced when more (HhH)_2_ domains are present, suggesting that even a few domains are capable of binding DNA (Belova et al., 2002; Rajan et al., 2010). Surrounding DNA is not necessary for activity, but it clearly enhances it. Furthermore, it has been shown that the topoisomerase V (HhH)_2_ domains can be used to enhance processivity of other enzymes (Pavlov, Belova, Kozyavkin, & Slesarev, 2002). Thus, the (HhH)_2_ domains may serve a similar role as sliding clamps that surround DNA (Hedglin, Kumar, & Benkovic, 2013), which enhance processivity by keeping polymerases closely associated with DNA while allowing movement along DNA. Unlike sliding clamps, which form a ring around DNA and need a clamp loader to assemble, the topoisomerase V (HhH)_2_ domains surround the DNA in a more extended manner and load into DNA without assistance.

The active site of topoisomerase V had not been observed in an accessible conformation before. All other known structures (Rajan et al., 2016; Rajan et al., 2013; Rajan et al., 2010; Taneja et al., 2006), aside from the structure of the isolated topoisomerase domain (Rajan et al., 2010), showed the active site obstructed by the (HhH)_2_ domains. Whereas it was clear that the (HhH)_2_ domains had to move to allow access to the active site, it was not clear how this is accomplished. The structure of the symmetric complex with DNA shows that the topoisomerase domain moves relative to the (HhH)_2_ domains through a single conformational change, a kink in the helix linking the first (HhH)_2_ domain and the topoisomerase domain. The domains themselves move as rigid bodies; the only change occurs in the linker helix. This simple mechanism allows the domains to separate and exposes the active site. In addition, the linker helix interacts with the DNA directly, suggesting that the conformational change could be triggered by protein DNA interactions. The movement of the domains also exposes a large, positively charged patch in the topoisomerase domain where the DNA enters the active site. In order to enter the active site, the DNA bends at two positions. There is small bend adjacent to the abasic site and a much larger bend where the DNA enters the topoisomerase domain. It is unlikely that the abasic site-induced bending reflects a feature of DNA binding and recognition as non-damaged DNA is easily relaxed by topoisomerase V. Surprisingly, the DNA is highly bent where it enters the topoisomerase domain and the bending has an unusual consequence, the double stranded DNA melts to allow only one strand to enter the active site. The two DNA strands separate, breaking the base pairing between them. One strand, the non-cleaved one, is anchored to the protein by clamping on a nucleobase. The other strand enters the active site and approaches the active site tyrosine. Both DNA strands exit the protein almost parallel to each other, suggesting that base pairing would resume once the DNA exits the protein.

The residues forming the topoisomerase active site were recognized based on structural and biochemical observations (Rajan et al., 2014; Taneja et al., 2006). The complex structure confirms that Arg131, Arg143, His200, Glu215, Lys218 and Tyr226 are involved in interacting with DNA and helps understand their role in the DNA cleavage/religation reaction. The two arginines, Arg131 and Arg144, interact directly with the phosphate group adjacent to the scissile bond (P-1) and are likely to interact also with the scissile bond phosphate (P0). Arg144 also forms a hydrogen bond to the OH group of the active site tyrosine. Mutations in either arginine led to very reduced activity (Rajan et al., 2014), suggesting an important role for these residues in the reaction. It was suggested (Rajan et al., 2014) that these arginines play a role in transition state stabilization, which would be broadly consistent with the observations from the complex structure. Glu215, an unusual residue due to its positive charge, comes close to the phosphate backbone and hydrogen bonds to His200. It was observed before that Glu215 reduces the binding affinity of the protein for DNA, which would be consistent with a negatively charged residue approaching the negatively charged phosphate backbone (Rajan et al., 2014). In the complex structure, His200 does not seem to interact directly with DNA so its role in the cleavage/religation reaction is not clear. Finally, the role of Lys218 is also not clear from the structure, as it is not observed in a position to interact with DNA, probably due to the absence of P0. It appears that it could interact with the phosphotyrosine intermediate helping stabilize it after cleavage; mutational studies show that the lysine is essential (Rajan et al., 2014). The combined structural and biochemical studies do suggest that the mechanism of cleavage and religation is different from the one employed by type IB enzymes. In type IB enzymes the histidine in the catalytic pentad interacts directly with DNA and the lysine plays a role in proton transfer. This does not appear to be the case in type IC topoisomerases. The presence of the glutamate in close contact with the phosphate backbone as well as the potential different role of His200 and Lys218 suggest that a distinct catalytic strategy is employed by topoisomerase V. Additional structural studies to capture covalent intermediates are needed to understand the cleavage/religation reaction.

Both type IB and IC topoisomerases use a swiveling mechanism to relax DNA. Structures of type IB enzymes in complex with DNA (Perry, Hwang, Bushman, & Van Duyne, 2006; Redinbo, Stewart, Kuhn, Champoux, & Hol, 1998) show that the DNA remains double helical and largely unbent and is surrounded by the protein (**Figure 5**). Cleavage and formation of a transient phosphotyrosine intermediate allows the intact strand to rotate until the broken free DNA strand is recaptured and ligated. Interaction of the DNA with the protein causes friction, which modulates the reaction (Koster et al., 2005). Single molecule experiments on topoisomerase V show a similar overall mechanism where friction also plays a role (Taneja et al., 2007), suggesting that type IB and IC enzymes employ the same general strategy. Given these parallel strategies, the presence of a sharp bend and a single stranded region in the complex of topoisomerase V was unexpected as type IB topoisomerases do not bend or melt the DNA (Perry et al., 2006; Redinbo et al., 1998). Single stranded DNA binding and recognition is part of the type IA mechanism, but these enzymes work by an enzyme bridged strand passage mechanism, which is fundamentally different from the swiveling mechanism employed by type IB and IC enzymes. These observations suggest that despite the apparent similarities, type IC enzymes employ a different relaxation strategy. Unlike type IB enzymes, type IC molecules bend the DNA to create a single stranded region that is likely to facilitate swiveling by freeing the two DNA strands around the cleavage site. It is not clear whether the rotation of the strands involves only rotation of the strands or also movement of the topoisomerase domain; it is possible that the topoisomerase domain moves as the strands rotate (**Figure 5**). Similar to type IB enzymes, the broken strand is captured after swiveling around the other strand. Finally, it is interesting to note that whereas type IB enzymes surround the DNA during the reaction, type IC enzymes do not appear to do so. The (HhH)_2_ domains surround the DNA, but their role seems to be to act as a processivity factor as these domains are not required for DNA relaxation (Rajan et al., 2010). Thus, while there are some overall similarities between the two subtypes, there are important differences that suggest that topoisomerase V employs a completely different relaxation mechanism.

**Figure 5.**
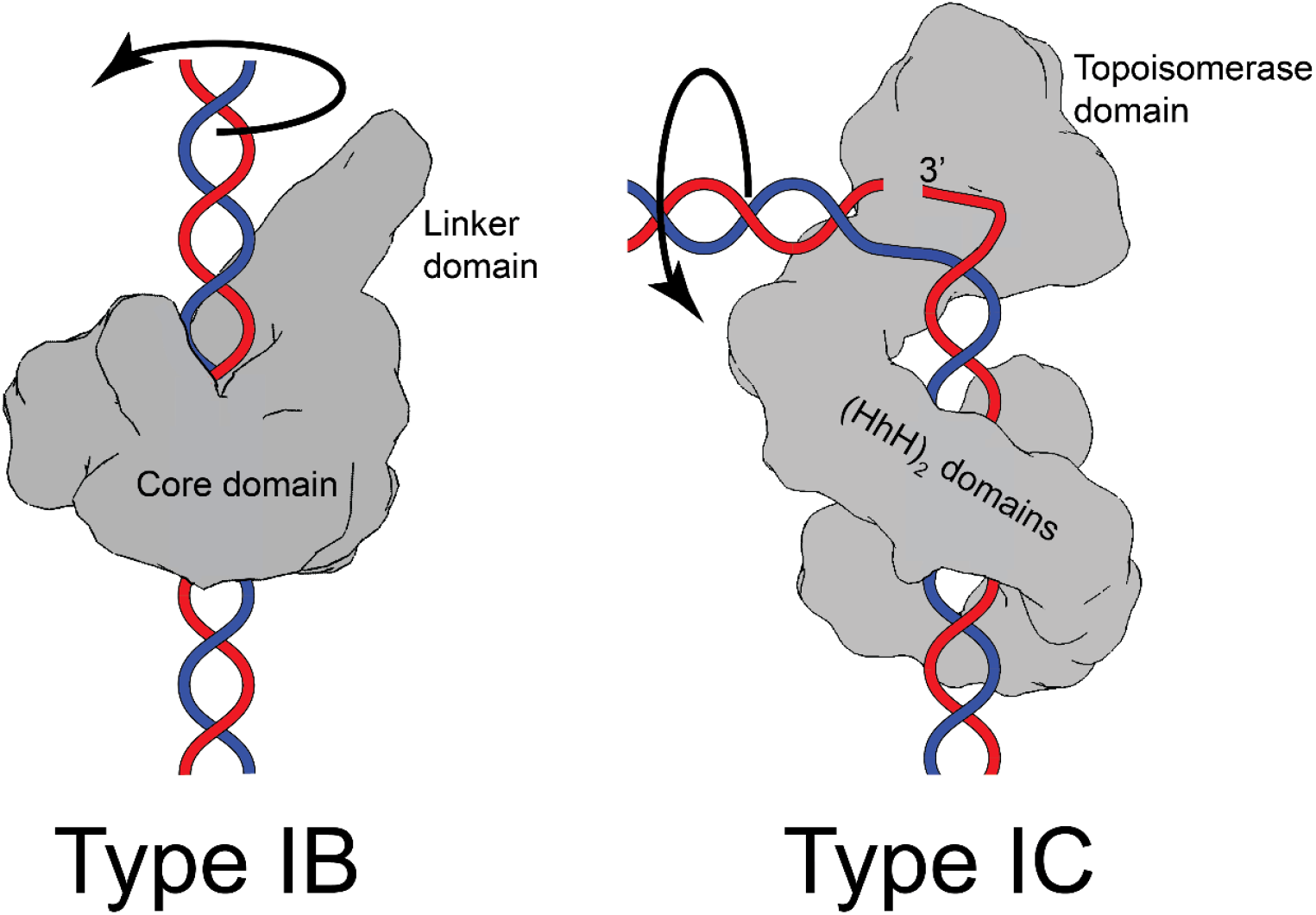
Comparison between the proposed mechanism of relaxation for type IB and type IC topoisomerases. Cartoons of the proposed relaxation mechanism for type IB enzymes, illustrated by a human topoisomerase complex, and topoisomerase V, a type IC enzyme. The proteins are shown as a surface based on their structure whereas the DNA is shown in cartoon form. **Left)** Type IB enzymes relax DNA by enclosing it and cleaving one strand, allowing for strand rotation or swiveling. The DNA is not bent or distorted. **Right)** Type IC topoisomerases do not surround the DNA around the active site region. They surround the DNA using tandem (HhH)_2_ domains, which appear to serve as a processivity factor. The DNA is highly bent around the active site region and this bending melts the two strands, allowing one of them to enter the active site. The way the two proteins interact with DNA is very different in both types. In addition, their DNA cleavage and religation mechanism appear to be different. In both cases, interactions between the protein and DNA create friction, which modulates the rate of the reaction. Supercoiling of the DNA creates torque, which drives the reaction. The type IB diagram was drawn based on the structure of human topoisomerase I in complex with DNA (PDB 1K4T) (Staker et al., 2002).

The structures presented here add significantly to our understanding of type IC enzymes and their mechanism. The structures show that an important role of the (HhH)_2_ domains is to surround the DNA and, in this way, keep the enzyme bound to DNA, probably allowing it to travel along DNA, and in this manner serve as a processivity factor. It is not clear from the structures how DNA lesions are recognized, but if the protein slides along the DNA it could scan it for lesions using the (HhH)_2_ domains containing the repair active sites. The topoisomerase domain is likely to be obstructed in the absence of DNA/protein interactions, but a simple conformational change around a linker helix exposes the active site once the protein is bound to DNA. The DNA bends tightly as it interacts with the protein around the active site region, leading to single stranded DNA formation, which may help facilitate strand rotation. Finally, the catalytic mechanism of DNA cleavage and religation appears to be different from the one employed by other topoisomerases, despite some very general similarities. All these observations indicate that topoisomerase V is a multifaceted enzyme that encompasses in the same polypeptide a novel topoisomerase domain, DNA repair domains, and a DNA clamp and that type IC enzymes are fundamentally different from all other topoisomerases. The structural, biochemical, and biophysical data help to establish firmly type IC topoisomerases as a completely different topoisomerase subtype with few sequence, structural, or mechanistic similarities to all the other subtypes.

## MATERIALS AND METHODS

### Protein purification

A fragment of Topo-V corresponding to residues 1 – 854 (Topo-97) with K809, K820, K831, K835, K846, and K851 mutated to alanine to remove the second AP lyase site (Topo-97(ΔRS2)) has been previously described (Rajan et al., 2016). For protein purification, Topo-97(ΔRS2) was transformed into *E. coli* BL21 Rosetta (DE3) cells. Protein induction and purification was done according to previously described protocols (Rajan et al., 2013; Rajan et al., 2010). Pure protein was concentrated to 55.6 mg/ml and stored in 50 mM Tris pH 8, 500 mM NaCl, and 1 mM DTT. Protein purity was assessed by polyacrylamide gel electrophoresis.

### Oligonucleotides

Oligonucleotides were purchased from Integrated DNA Technologies (IDT, Coralville IA) at 250 nmol. The sequences of the oligonucleotides used are shown below.

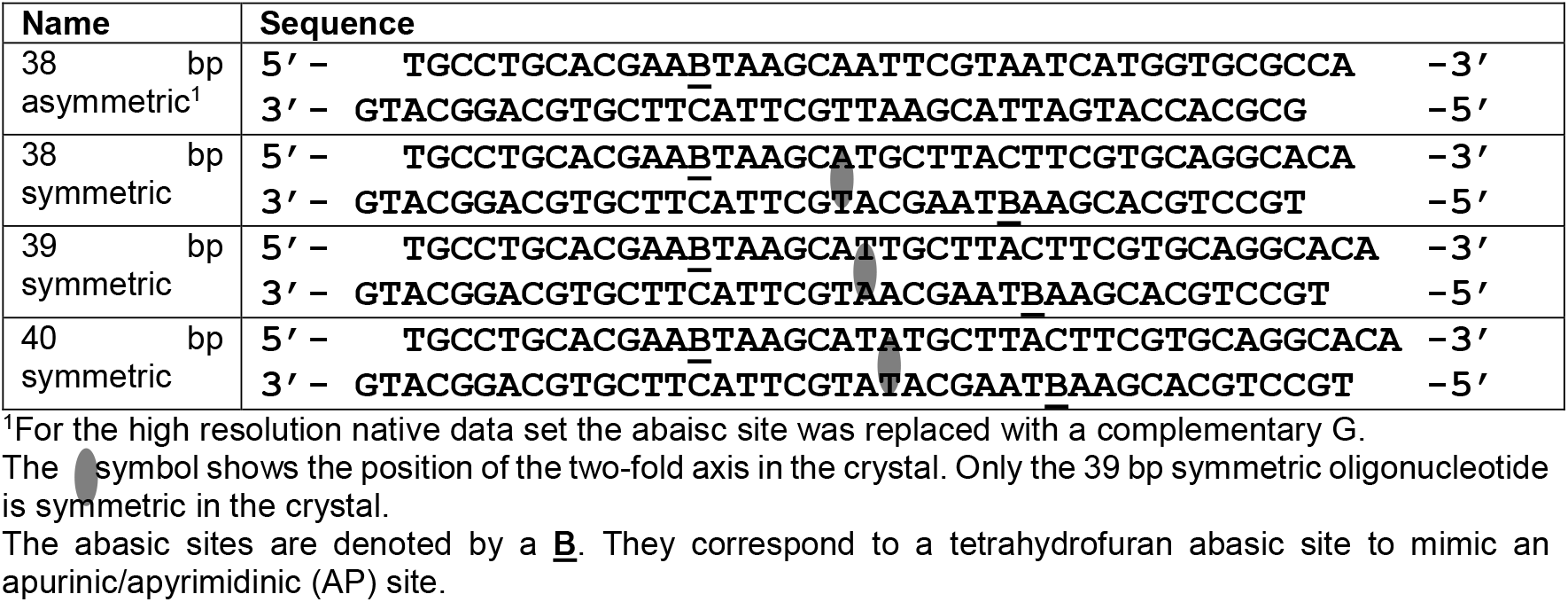

Individual oligonucleotides were resuspended to 1mM in water. For annealing, complementary oligonucleotides were mixed at an equimolar ratio, heated to 85°C for 2.5 minutes, cooled down to 5°C below their calculated melting temperature for 5 minutes, and then transferred to ice for at least 10 minutes. Annealed oligonucleotides were used directly in crystallization experiments.

### Crystallization, data collection, and structure determination

For all crystallization experiments, Topo-97(ΔRS2) was mixed with the corresponding oligonucleotide using a stoichiometric ratio of 1.25:1 DNA to protein in 1X DNA binding buffer. Reactions were incubated for thirty minutes at 65° C. For the asymmetric DNA experiments, molar ratios of 40:32 µM and 60:48 µM DNA to protein were used whereas for the symmetric DNA experiments a ratio of 60:48 µM DNA to protein yielded the best crystals. 5X DNA binding buffer was: 250 mM sodium acetate pH 5.0 at 65°C, 150 mM sodium chloride, 5 mM magnesium chloride.

All crystallizations were done by vapor diffusion at 30° C in a hanging drop setup. Topo-97(ΔRS2) with asymmetric DNA crystals typically started to appear three days after setup and continued to appear for up to two weeks. For these crystals, the well solution was added in 1:1 or 2:1 complex to well solution ratio; it was found that the 2:1 ratio would typically give fewer and larger crystals in the drops. Topo-97(ΔRS2) with symmetric DNA crystals started to appear within minutes of setting up the trays in 1:1 or 2:1 well to complex ratio. Subsequently it was discovered that adding phosphotungstic acid (H_3_PW_12_O_40_) at between 12.5 – 15 µM made the crystals grow slower and larger and the morphology changed from plates to box-like crystals. Exact conditions for the crystallization experiments are shown in **Table I**.

**Table I.**
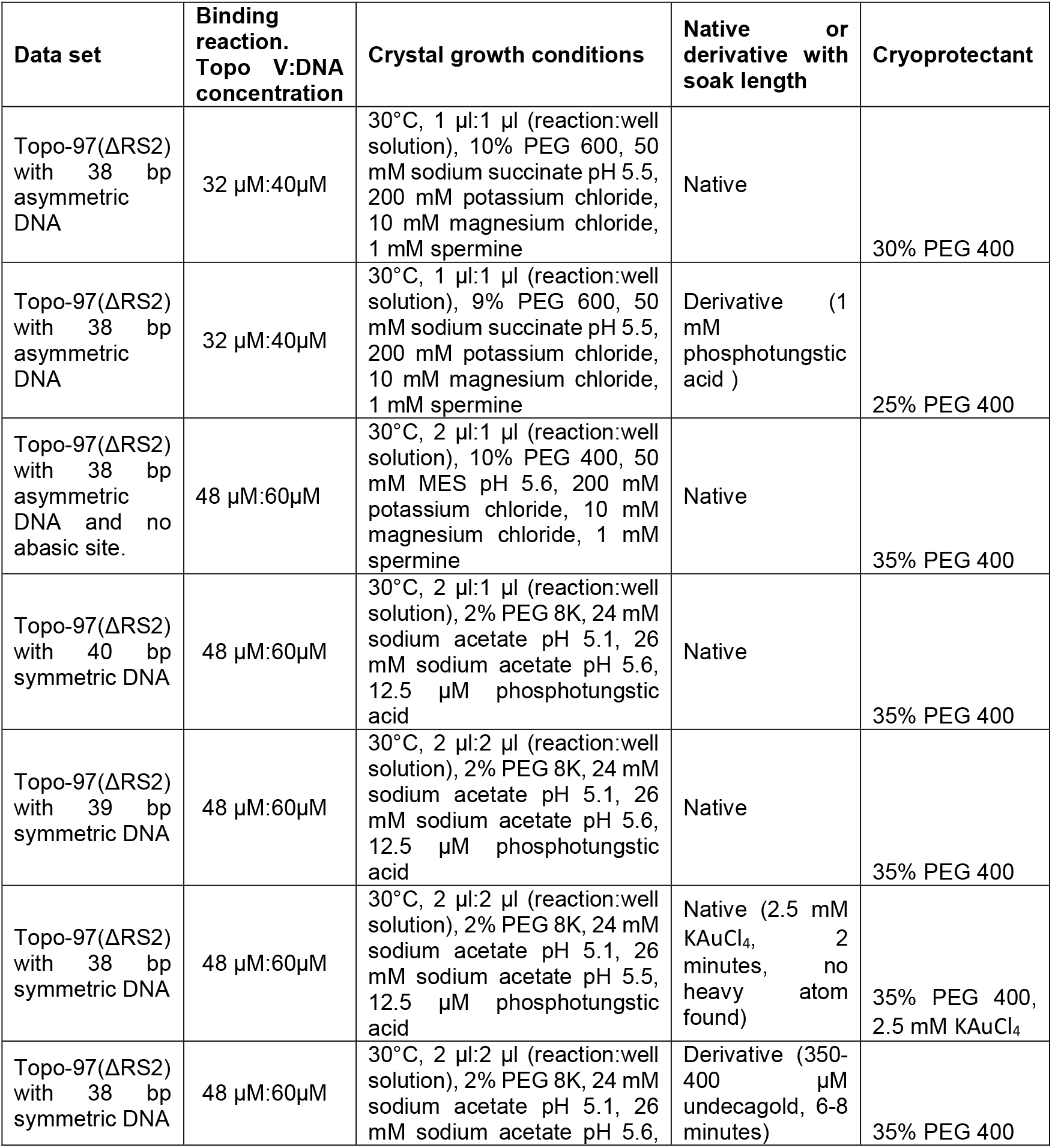
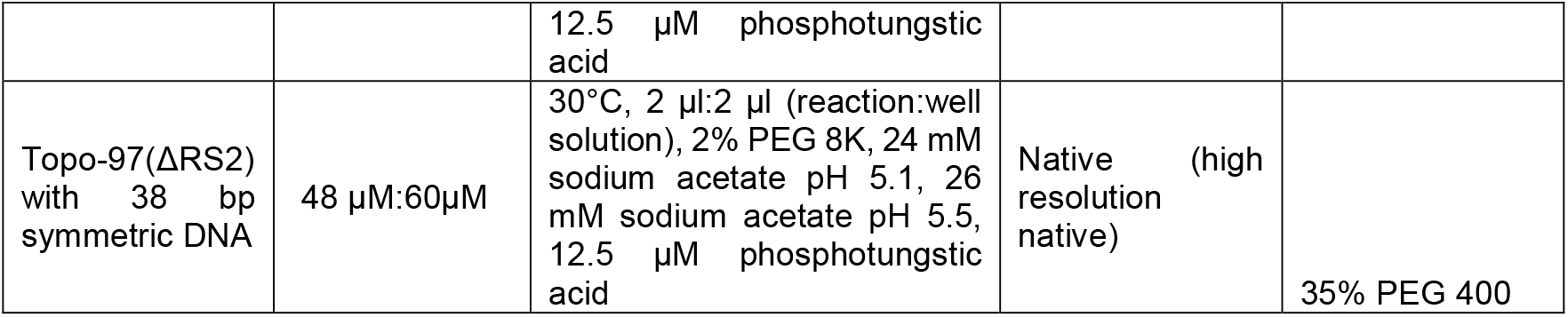
Crystallization conditions.

For data collection, crystals were first transferred to cryoprotectant (see **Table I**) and then flash frozen in liquid nitrogen. All diffraction data were collected at the Life Science Collaborative Access Team station (LS CAT) at the Advanced Photon Source (APS) in Argonne National Laboratory. All data were processed using XDS (Kabsch, 1993), aimless (Evans & Murshudov, 2013), autoPROC (Vonrhein et al., 2011), and other programs of the CCP4 suite (Winn et al., 2011). Data anisotropy was analyzed with the StarAniso server (Tickle et al., 2018) or with autoPROC (Vonrhein et al., 2011).

The structure of Topo-97(ΔRS2) with an asymmetric 38 bp DNA was solved by a combination of Molecular Replacement and heavy atom phasing. A weak Molecular Replacement solution with Phaser (McCoy et al., 2007) was found against a 6 Å data set using a 61 kDa fragment of topoisomerase V (Taneja et al., 2006). The Molecular Replacement solution did not show any additional protein regions or DNA. A phosphotungstic acid derivative was prepared by soaking crystals in 1 mM phosphotungstic acid for 2 minutes before flash freezing them. The Molecular Replacement model was used to locate the heavy atoms. Phasing was done with SHARP (Vonrhein, Blanc, Roversi, & Bricogne, 2007) (**Table II**). The map, even at 6 Å resolution, show the position of two protein monomers and two double stranded DNA oligonucleotides. The model of a 97 kDa fragment of topoisomerase V (Rajan et al., 2016) was used to build most of the protein and the DNA molecules were built starting from idealized DNA. Subsequently, a higher resolution, anisotropic data set with DNA without the abasic site was used to refine the structure to 3.24 Å resolution in the best direction using Buster (Roversi, Irwin, & Bricogne, 1996) and Phenix (Adams et al., 2010). Manual model building was done using Coot (Emsley & Cowtan, 2004; Emsley, Lohkamp, Scott, & Cowtan, 2010). The final model consists of two protein monomers, one full DNA molecule, and two half-length DNA molecules. The full DNA molecule, which spans 42 nucleotides per strand with 40 of them forming basepairs, binds both protein monomers while the other two DNA molecules bind one protein monomer each and sit on crystallographic axes generating crystallographic dimers made of symmetric full-length DNA molecules each with two protein monomers. The two protein complexes are not identical, as the DNA sits differently in each one. The final R_work_ and R_free_ for the model are 23.35% and 26.38%, respectively. The model has excellent stereochemistry. Molprobity (Davis, Murray, Richardson, & Richardson, 2004) shows that 97.22% of the residues in the Ramachandran plot are in favored regions and no residues are in disallowed regions. The model has a root mean square deviation (rmsd) of 0.004 Å for bond lengths and 0.54° for bond angles (**Table IV**).

**Table II.**
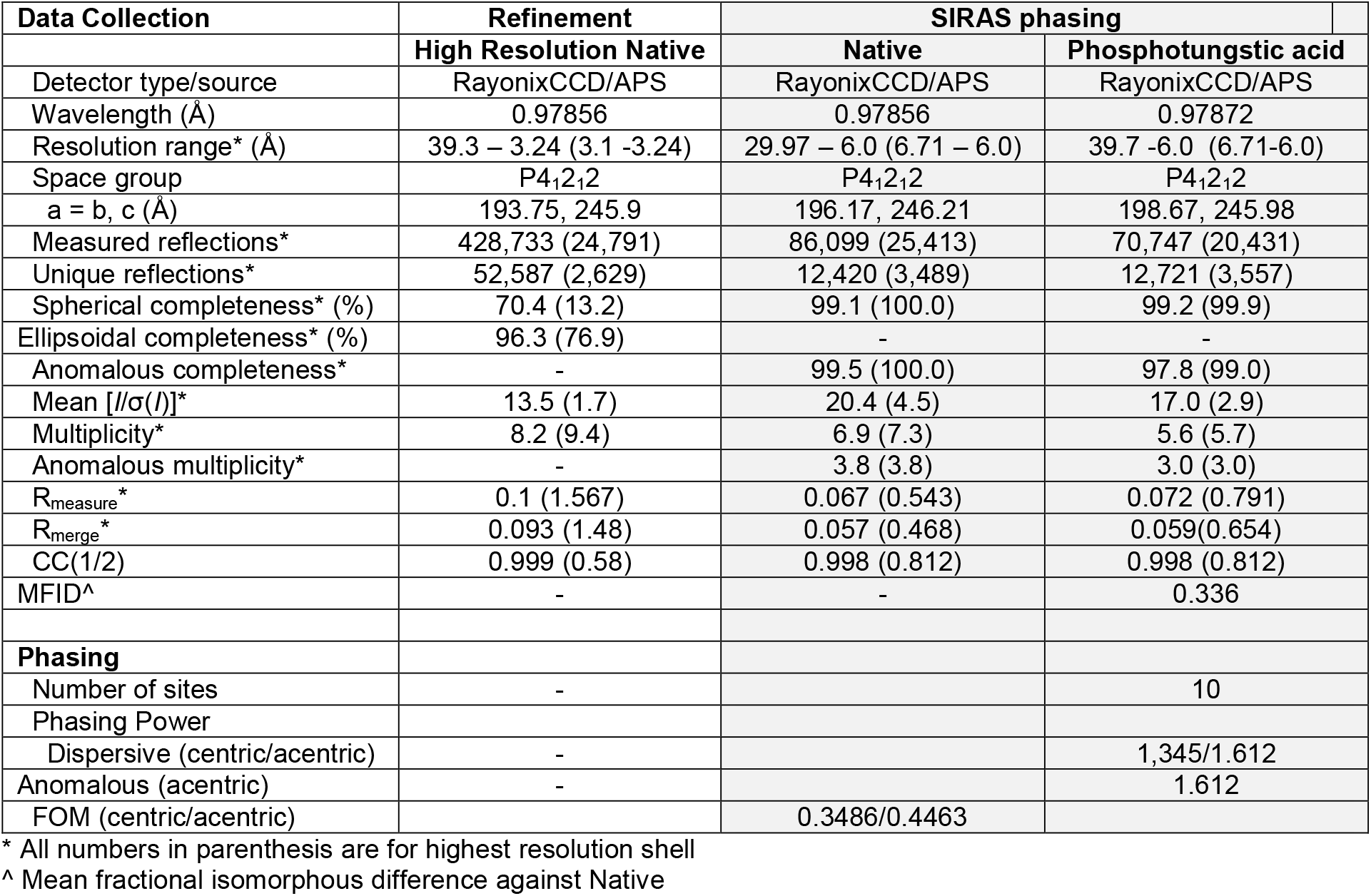
Data collection and phasing statistics for asymmetric complex.

The structure of Topo-97(ΔRS2) with a symmetric 38 bp DNA was solved by heavy atom phasing using a 0.8 nm positively charged undecagold (Nanoprobes, Inc., Yaphank NY) derivative (**Table III**). The undecagold derivative was prepared by soaking crystals in 350-400 µM undecagold for 6-8 minutes before flash freezing them. Data from 5 different crystals were merged using Blend (Foadi et al., 2013) to create a single derivative data set. Phenix (Adams et al., 2010) was used to locate a single heavy atom. Phasing was done with SHARP (Vonrhein et al., 2007) treating the undecagold as a spherically averaged cluster. The electron density map showed two protein monomers bound to a single 38 bp oligonucleotide. The model of Topo-97(ΔRS2) in complex with an asymmetric 38 bp DNA was used to build the two protein monomers whereas the DNA was built starting from ideal DNA. The structures of Topo-97(ΔRS2) with symmetric 39 bp and 40 bp symmetric DNA were obtained by refining the Topo-97(ΔRS2) in complex with an asymmetric 38 bp DNA. In all cases, anisotropic data sets were used to refine the structures using Buster (Roversi et al., 1996) and Phenix (Adams et al., 2010). Manual model building was done using Coot (Emsley & Cowtan, 2004; Emsley et al., 2010). Unlike the asymmetric complex, the symmetric complexes show the position of the abasic sites. In each case, the final model consists of two protein monomers and one full DNA molecule. Whereas the conformation of the protein is very similar in all cases, the DNA shows different distortions around the abasic sites. The final R_work_/R_free_ for the Topo-97(ΔRS2) with symmetric 38 bp, 39 bp and 40 bp DNA models are 25.71%/28.77%, 21.84%/25.11%, and 22.89%/27.00%, respectively. All models have excellent stereochemistry. **Figure 4 – Figure Supplement 1** shows density around the active site region. Molprobity (Davis et al., 2004) shows that 96%, 95.8%, and 97.5% of the residues in the Ramachandran plot are in favored regions and no residues are in disallowed regions for the 38 bp, 39 bp, 40 bp DNA models. The models have a root mean square deviation (rmsd) of 0.002 Å for bond lengths and 0.42-0.49° for bond angles (**Table IV**).

**Table III.**
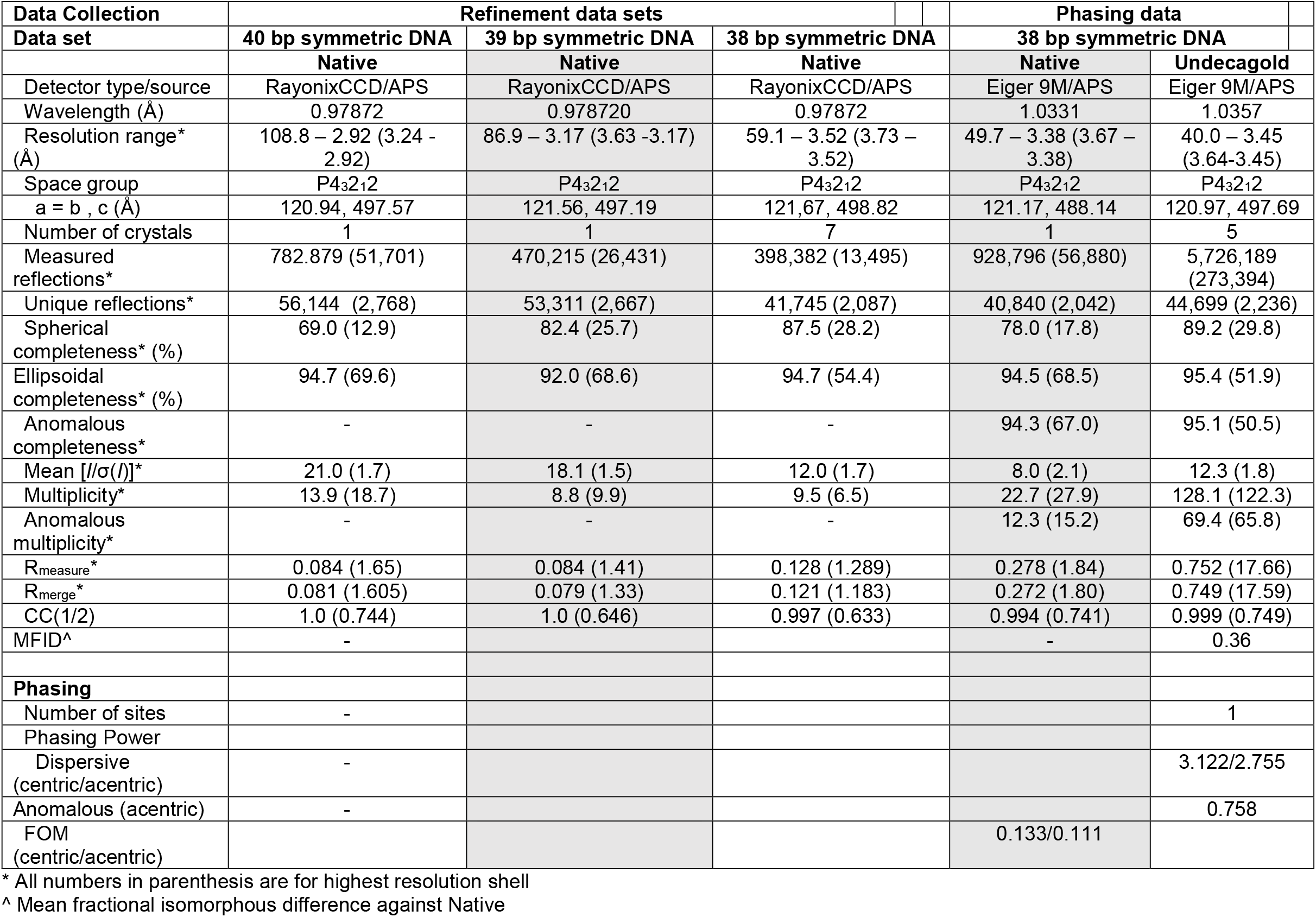
Data collection and phasing statistics for symmetric complexes.

**Table IV.** Refinement statistics for all complexes.

| Refinement | 40 bp<br>symmetric<br>DNA | 39 bp<br>symmetric<br>DNA | 38 bp<br>symmetric<br>DNA | 38 bp asymmetric<br>DNA (no abasic<br>site) |
| --- | --- | --- | --- | --- |
| Resolution (Å) | 58.76 – 2.92 | 59.04 – 3.17 | 59-10 – 3.52 | 39.28 – 3.24 |
| Number of reflections<br>working/test<sup>a</sup> | 56,123/2,845 | 53,291/2,690 | 41,720/2,217 | 52,466/2,599 |
| *R* (working set; %)<sup>b</sup> | 25.71 (43.55) | 21.84 (34.04) | 22.29 (33.41) | 23.35 (39.48) |
| *R*<sub>free</sub> (test set; %)<sup>c</sup> | 28.77 (26.08) | 25.11 (69.52) | 27.00 (38.76) | 26.38 (25.78) |
| Structure Quality |  |  |  |  |
| Protein atoms | 13,614 | 13,604 | 13,604 | 13,014 |
| DNA atoms | 1,606 | 1,619 | 1,561 | 2,868 |
| Other atoms | 32 | 4 | 5 | 44 |
| RMS deviations in bond<br>lengths (Å) | 0.002 | 0.002 | 0.002 | 0.004 |
| RMS deviations in bond<br>angles (°) | 0.489 | 0.442 | 0.421 | 0.540 |
| Average B factor (Å<sup>2</sup>) (DNA) | 110.87 | 117.99 | 138.26 | 191.74 |
| Average B factor (Å<sup>2</sup>) (protein) | 97.13 | 104.43 | 129.73 | 135.94 |
| Ramachandran plot<sup>d</sup> |  |  |  |  |
| Favored regions (%) | 97.52 | 95.81 | 95.99 | 97.22 |
| Outliers (%) | 0.24 | 0.41 | 0.41 | 0 |
| Distribution Z-score | -1.99 | -1.90 | -1.27 | -1.51 |
| Clashscore | 5.93 | 4.76 | 4.94 | 4.86 |
| Molprobity score | 1.42 | 1.53 | 1.53 | 1.4 |

<sup>a</sup>Numbers in parenthesis correspond to highest resolution shell.
<sup>b</sup> $R_{\text{work}} = \sum ||F_o| - |F_c|| / \sum |F_o|$ , where $|F_o|$ is the observed structure factor amplitude and $|F_c|$ the calculated structure factor amplitude.
<sup>c</sup> $R_{\text{free}} = R_{\text{factor}}$ based on 5% of the data excluded from refinement.
<sup>d</sup>As reported by Molprobity (Davis et al., 2004).

### Figures

Figures for the atomic models were created using Pymol (DeLano, 2002) and Chimera (Pettersen et al., 2004). Electrostatic surfaces were calculated using the program APBS (Baker, Sept, Joseph, Holst, & McCammon, 2001).

### Accession numbers

Atomic coordinates and structure factors for the reported crystal structures have been deposited with the Protein Data Bank under the accession numbers XX1, XX2, XX3, XX4.

## ACKNOWLEDGEMENTS

We thank E. Smith and V. Tokars for comments on the manuscript and members of the Mondragón laboratory for help and suggestions. Research was supported by the NIH (R35-GM118108 to A.M.). We acknowledge the help from the Northwestern University Structural Facility and the beamline scientists at LS-CAT/Sector 21 at the Advanced Photon Source, Argonne National Laboratory. LS-CAT/Sector 21 was supported by the Michigan Economic Development Corporation and the Michigan Technology Tri-Corridor. Support from the R.H. Lurie Comprehensive Cancer Center of Northwestern University to the Structural Biology Facility is acknowledged.

## Competing Financial Interests

The authors declare no competing financial interests.

**Figure 1 – Figure Supplement 1.**
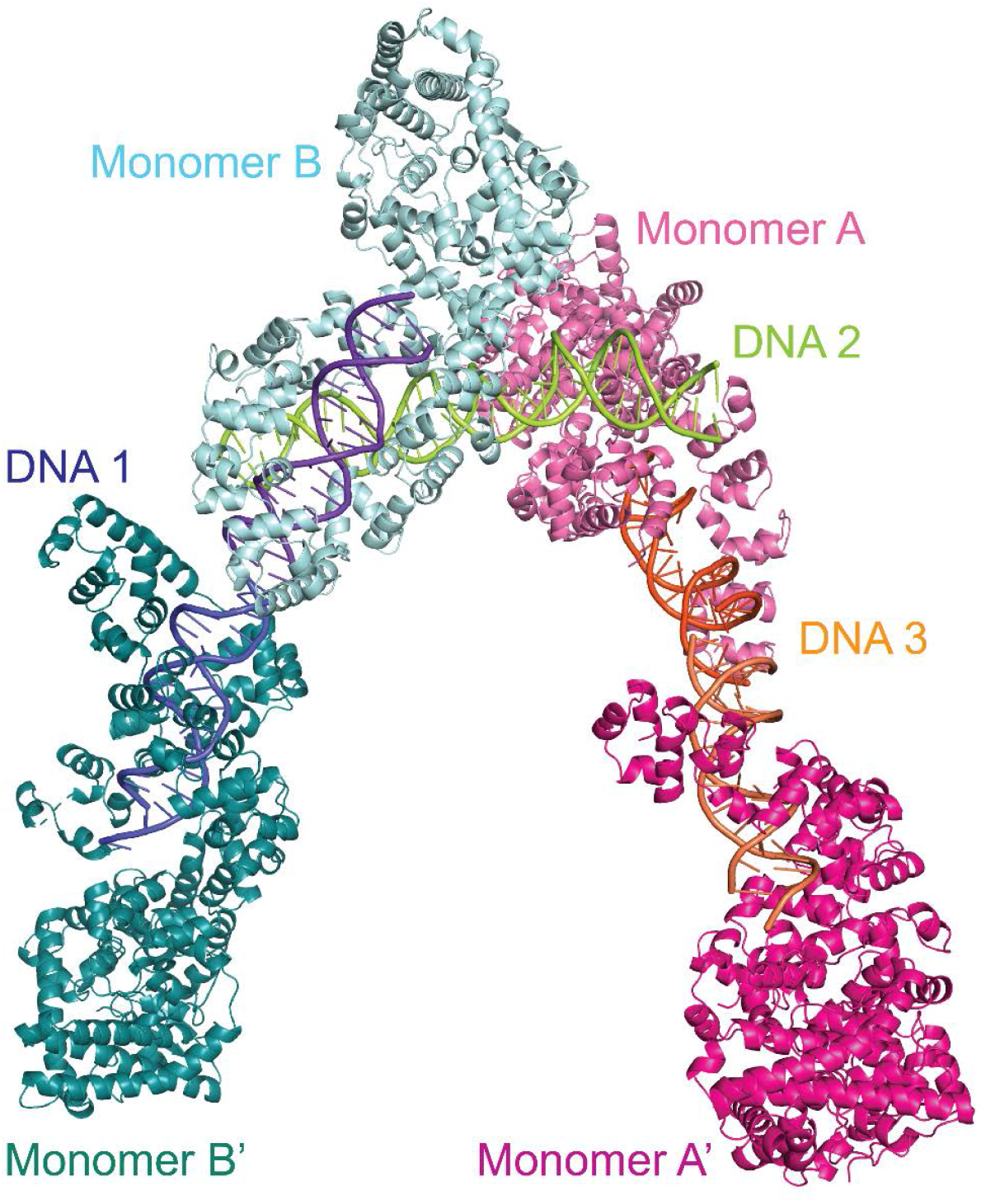
Dimers in the crystal of the topoisomerase V with asymmetric DNA complex. The diagram shows the way the molecules are arranged in the crystal. Two of the DNA molecules, DNA 1 and DNA 3, each bind two protein monomers (A, A’ and B, B’) each surrounding one half of the DNA. The center of the DNA in these two molecules sits on crystallographic two-fold axes. The third DNA molecule, DNA 2, binds two protein monomers (A and B), but in this case the DNA is not surrounded by the protein.

**Figure 2 – Figure Supplement 1.**
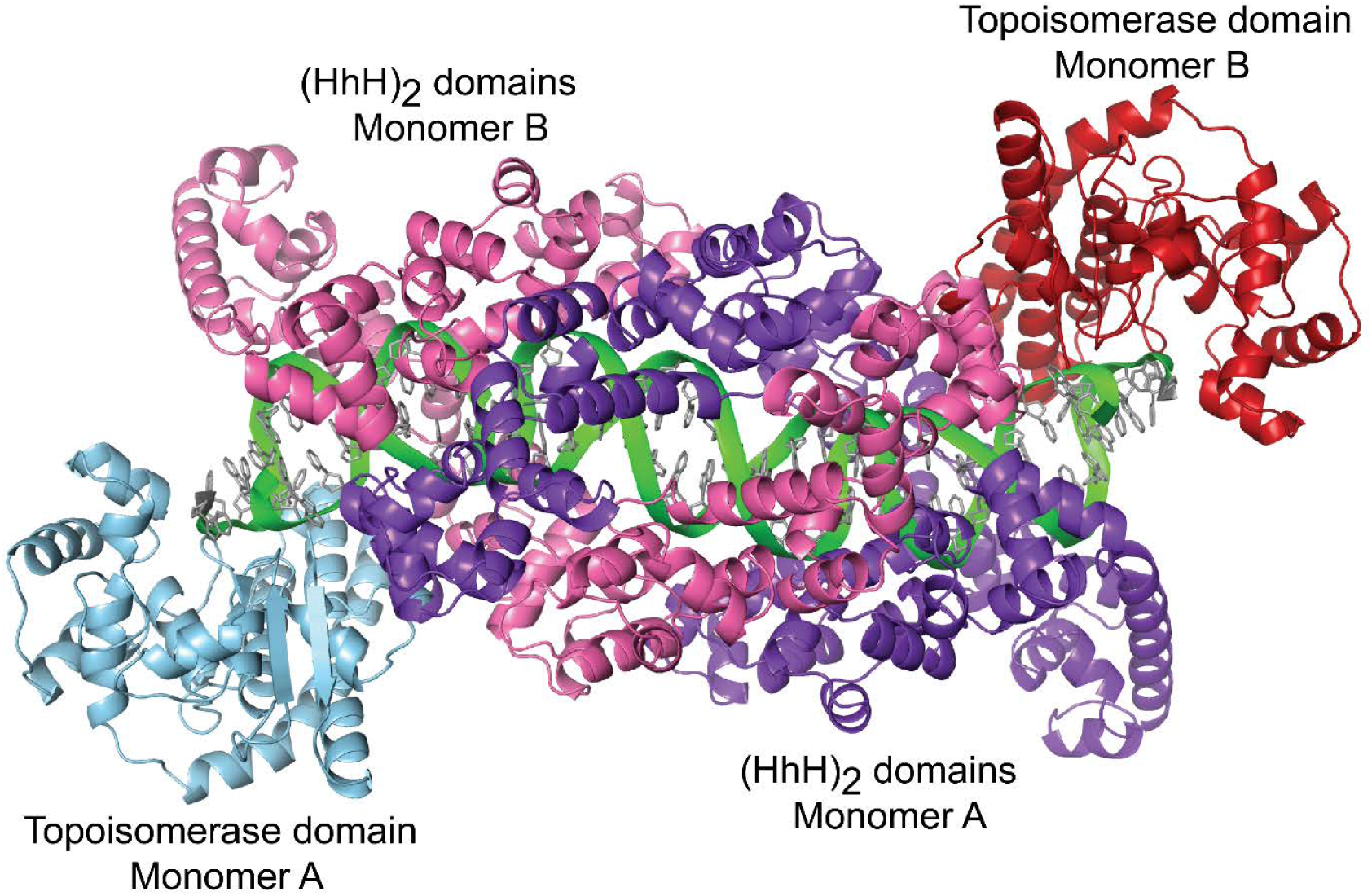
Dimer in the crystal of topoisomerase V with symmetric 39 bp DNA. The diagram shows the way the molecules are arranged in the crystal. Each dimer in the crystal is formed by one DNA molecule (green), and two protein monomers (blue and purple, and red and pink). For the 39bp DNA, the dimer is almost perfectly symmetric with the two halves of the DNA identical. This is not the case in the 38 bp and 40 bp symmetric DNA complexes, where the DNA is different around the abasic site in each half. Despite the small differences in the DNA, the protein is identical in all complexes.

**Figure 2 – Figure Supplement 2.**
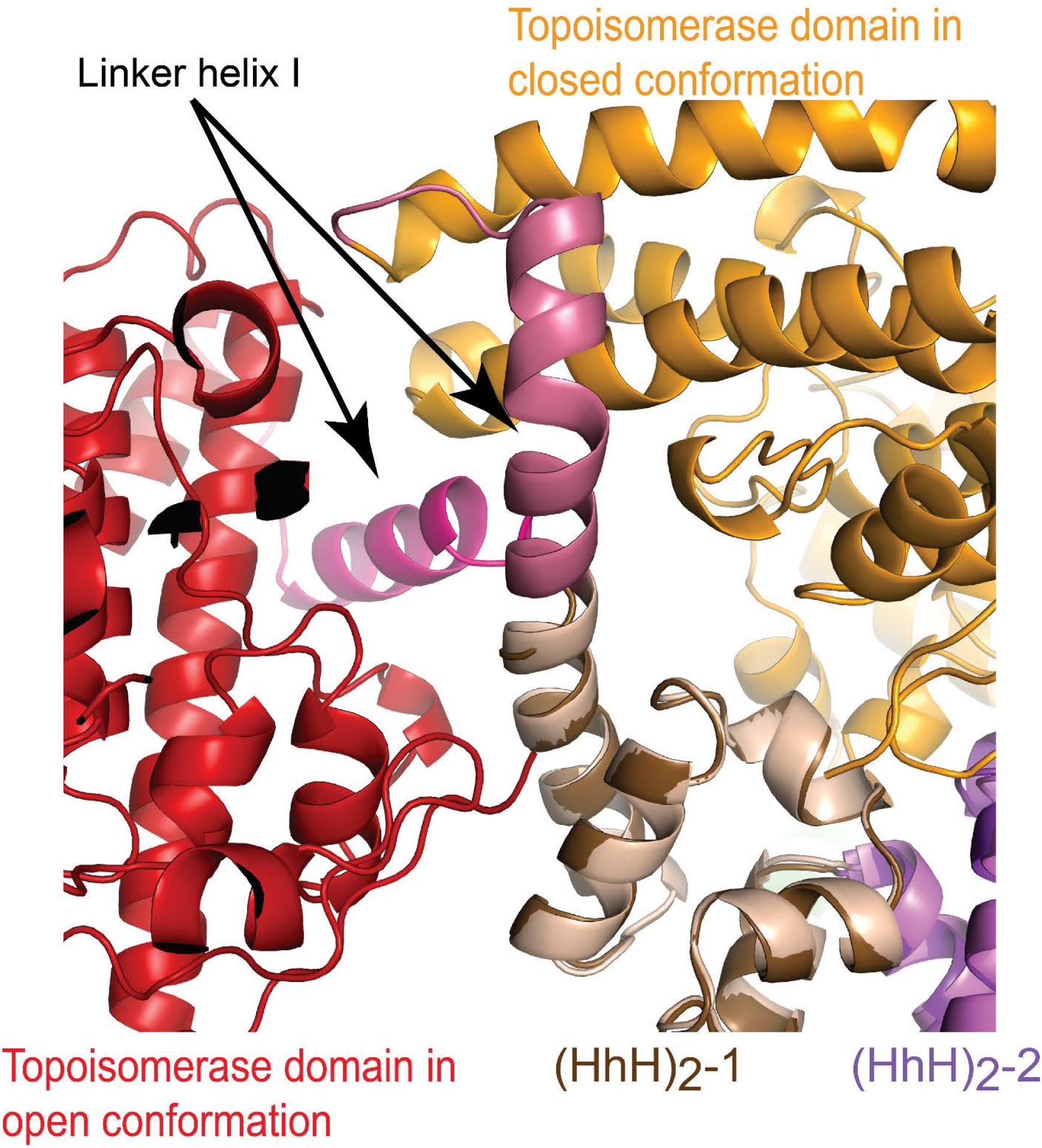
A conformational change in Linker helix I exposes the topoisomerase active site. Superposition of the closed and open conformations of topoisomerase V in complex with DNA. For the comparison, (HhH)_2_ domains 1 and 2 were superposed in the asymmetric DNA (closed) and symmetric DNA (open) complexes. In the open complex, the topoisomerase domain is colored red, whereas it is colored yellow in the closed complex. The only major change is the breaking of the linker helix (pink). This conformational change causes the topoisomerase domain to move away from the (HhH)_2_ domains and expose the active site.

**Figure 3 – Figure Supplement 1.**
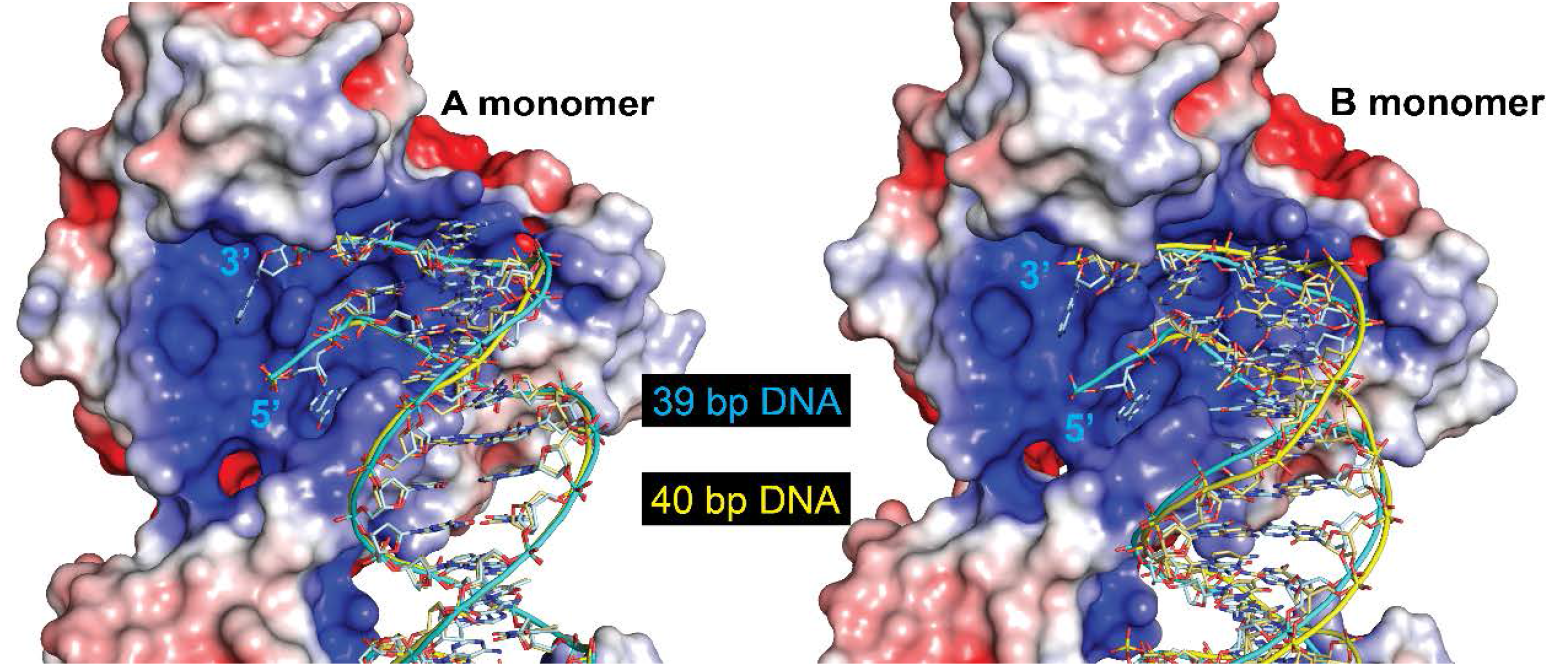
The protein can accommodate different length DNA entering the active site. The figure shows the DNA in the 39 bp symmetric DNA complex (cyan) with the DNA from the 40 bp DNA symmetric complex (yellow) superposed. The electrostatic surface corresponds to the 39 bp symmetric complex. The path of the DNA is identical in the A monomer (left), but follows a slightly different path in the B monomer (right). The change in path is needed to accommodate the different lengths of the two halves of the DNA. Even though the path is slightly different in the B monomer, the ends of the DNA are in the same position and the DNA enters the active site identically.

**Figure 4 – Figure Supplement 1.**
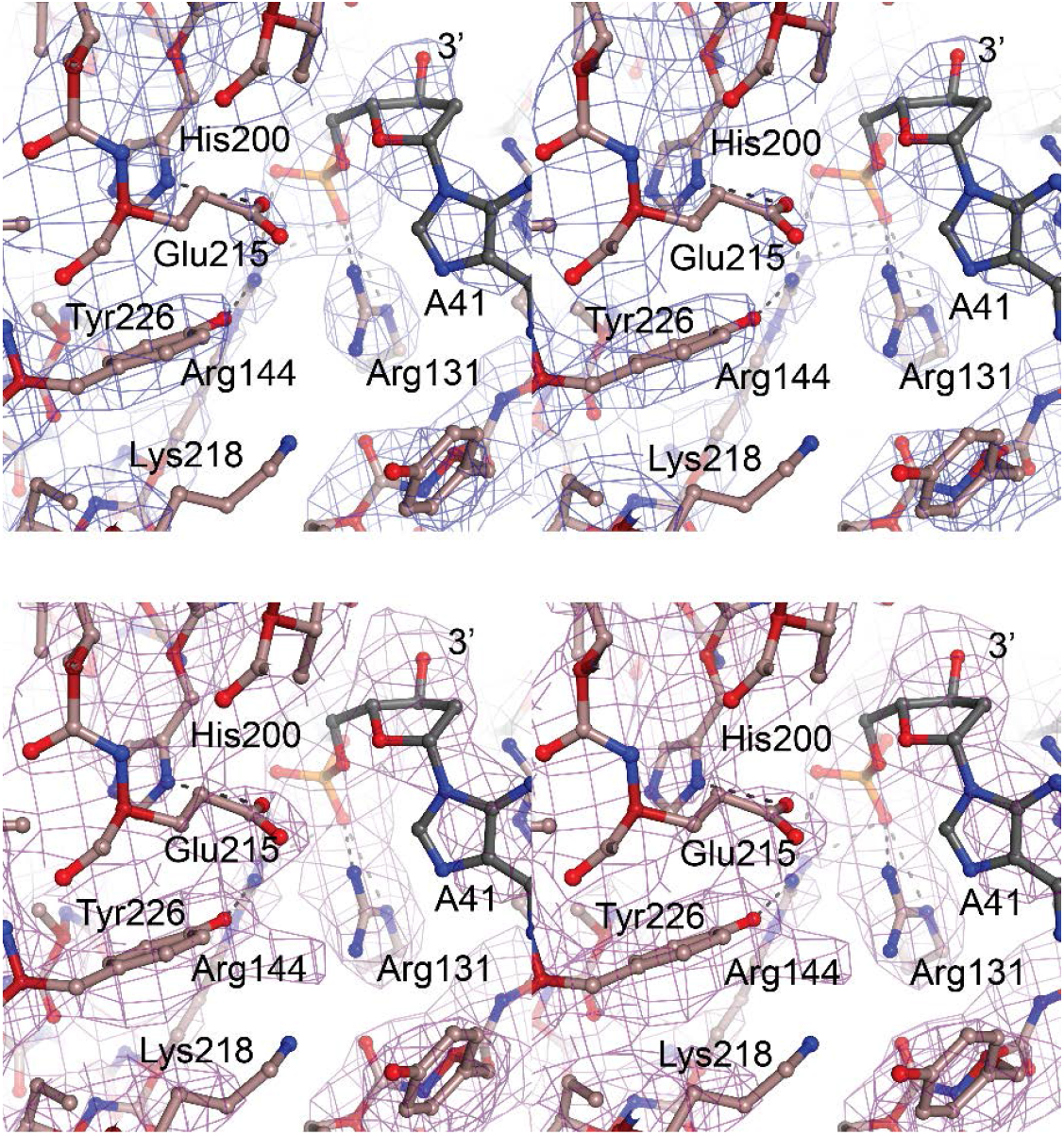
Electron density maps of the topoisomerase V in complex with 39 bp symmetric DNA. **A)** Stereo diagram of the active site region showing a 2mFo-DFc map. The map it is contoured at the 1.4 σ level. **B)** Stereo diagram of the same region as in **A**, but showing a simulated annealing omit map contoured at the 1.0 σ level. Both maps were calculated with Phenix (Adams et al., 2010).

## Notes

### Competing Interest Statement

The authors have declared no competing interest.

